# Identifying Temporal Correlations Between Natural Single-shot Videos and EEG Signals

**DOI:** 10.1101/2023.09.19.558394

**Authors:** Yuanyuan Yao, Axel Stebner, Tinne Tuytelaars, Simon Geirnaert, Alexander Bertrand

## Abstract

**Objective:** Electroencephalography (EEG) is a widely used technology for recording brain activity in brain-computer interface (BCI) research, where understanding the encoding-decoding relationship between stimuli and neural responses is a fundamental challenge. Recently, there is a growing interest in encoding-decoding natural stimuli in a single-trial setting, as opposed to traditional BCI literature where multi-trial presentations of synthetic stimuli are commonplace. While EEG responses to natural speech have been extensively studied, such stimulus-following EEG responses to natural video footage remain underexplored.

**Approach:** We collect a new EEG dataset with subjects passively viewing a film clip and extract a few video features that have been found to be temporally correlated with EEG signals. However, our analysis reveals that these correlations are mainly driven by shot cuts in the video. To avoid the confounds related to shot cuts, we construct another EEG dataset with natural single-shot videos as stimuli and propose a new set of object-based features.

**Main Results:** We demonstrate that previous video features lack robustness in capturing the coupling with EEG signals in the absence of shot cuts, and that the proposed object-based features exhibit significantly higher correlations. Furthermore, we show that the correlations obtained with these proposed features are not dominantly driven by eye movements. Additionally, we quantitatively verify the superiority of the proposed features in a match-mismatch (MM) task. Finally, we evaluate to what extent these proposed features explain the variance in coherent stimulus responses across subjects.

**Significance:** This work provides valuable insights into feature design for video-EEG analysis and paves the way for applications such as visual attention decoding.

## 1 Introduction

Electroencephalography (EEG) is a non-invasive technology to record the electrical activity of the brain through electrodes attached to the scalp. Due to its high temporal resolution, affordability and portability, EEG has found extensive applications in brain-computer interface (BCI) research. A key challenge in BCIs is understanding the encoding-decoding relationship between stimuli and EEG responses. Early approaches, e.g., the event-related potential (ERP) approach [1], heavily relied on short synthetic sensory stimuli, such as tone beeps or sudden visual events. Participants were repeatedly presented with the same synthetic stimulus, and the neural response was obtained by averaging the EEG signals across trials to deal with the low signal-to-noise ratio (SNR) of EEG. Using synthetic stimuli leads to more deterministic neural responses, such that averaging multiple EEG trials allows to remove the uncorrelated background noise while retaining the neural signals of interest. However, such multi-trial paradigms often lead to participant fatigue and are impractical for real-life applications involving natural continuous stimuli like audio and video footage that emerge in everyday life applications, in which case the stimuli are only presented once. Consequently, there has been a growing need to develop new paradigms that use natural stimuli in a single-trial context. The challenge lies in the fact that the stimulus-following responses are non-deterministic (as opposed to ERPs), and that they are buried under all kinds of EEG background noise, requiring more advanced signal processing tools to decode them, in particular since multi-trial averaging is not an option.

Therefore, when working with natural single-trial stimuli, two crucial elements are (1) a good representation of the stimulus that correlates well with the EEG and (2) an appropriate model that captures the relationship between the stimulus representation and the EEG response, while removing background EEG. Extensive research has been conducted on auditory-EEG analysis for natural speech, where various useful speech representations have been proposed, ranging from low-level features such as the spectrogram [2] and speech envelope [3, 4, 5], to high-level information such as phonemes [6] and semantic context [7]. The most common models are linear models, which can be roughly categorized into two groups: forward models and backward models. Linear forward models assume that the EEG signal consists of a stimulus response superimposed to background EEG, where the former is typically modelled as a convolution between a proper stimulus representation (e.g. a speech envelope) and a so-called temporal response function (TRF). The TRF can be estimated using, e.g., Least Squares (LS), and the EEG signals can be predicted from the audio features using the estimated TRF [3, 6, 7]. Backward models, on the other hand, reconstruct the stimulus as a linear combination of (lagged) EEG channels [4, 5]. A hybrid encoding-decoding model based on Canonical Correlation Analysis (CCA) was proposed in [8], where linear transformations were applied to both the speech envelope and the EEG signals such that the latent representations were maximally correlated. However, linear models have an inherent limitation in capturing the nonlinear dynamics of the brain. Moreover, using linear models also makes the results very dependent on the handcrafted feature engineering of the stimulus representation. Therefore, deep learning methods have been receiving increasing attention in recent years [9]. For instance, [10] decoded the auditory brain using deep learning-based CCA, and in [11], a Long Short-Term Memory-based model was proposed to discriminate whether a pair of an EEG segment and speech envelope correspond to each other or not. One direct application of audio-EEG analysis is auditory attention decoding, which has paved the way for technologies such as neural-steered hearing aids [12].

While there have been successful attempts to decode natural audio from EEG, the decoding of natural video footage from EEG has received less attention. The high-dimensional nature of the video signals poses challenges in finding useful representations. A possible approach is to not explicitly take the video stimulus into account in the modeling and extract common EEG components across the EEGs of multiple subjects watching the same video using methods such as Correlated Component Analysis (CorrCA) [13, 14, 15, 16]. By construction, the EEG responses that are coherent across subjects can only be time-locked to the visual stimuli since only the video stimuli is shared during all the EEG measurements. However, the link between the extracted EEG components and the video is unclear, making these components not very interpretable, in particular in regard to which features in the video drive the correlation. Alternatively, stimulus-aware algorithms such as CCA can be used to analyze the encoding-decoding relationship between the (specific features of) video stimulus and individual EEG signals. In this case, the design of relevant (low-dimensional) video features becomes crucial. In previous studies, the mean velocity of pixels calculated from the optical flow and the mean temporal derivative of pixel intensity (temporal contrast) were shown to be correlated with individual EEG signals, suggesting that they could be visual features that elicit strong EEG responses [17, 18].

In this work, we argue that shot cuts, which refer to sudden changes in the camera viewpoint or scene, have a significant impact on stimulus-aware video-EEG analysis, using previously proposed video features, as well as on (stimulus-unaware) multi-subject EEG analysis. Moreover, we recorded a new EEG dataset with subjects watching a set of single-shot videos containing a single moving object (i.e., a person). These single-shot single-object videos were specifically chosen to avoid introducing confounds related to shot cuts and to reduce the complexity of the stimuli. We demonstrate that the previously proposed mean velocity of pixels and temporal contrast are not relevant enough to generate significant correlations in the absence of shot cuts, and propose new object-based versions of these features that are more relevant, leading to significantly higher correlations. Moreover, we show that the EEG components obtained with the new features are not dominantly driven by eye movements, which are usually considered confounds. We further demonstrate that the proposed features are of better quality by their lower error rates in a match-mismatch (MM) task. Finally, we perform a multi-subject EEG analysis and calculate the proportion of variance in the coherent stimulus responses explained by the proposed features.

The structure of the paper is outlined as follows: In Section 2, we review the mathematical tools that we used in our analysis and introduce the proposed video features. Section 3 describes the experimental protocol in detail. The results are presented in Section 4, followed by further discussions in Section 5. Finally, in Section 6, we draw conclusions based on our findings.

## 2 Methods

To quantify and identify the temporal coupling between signals, we choose CCA in the context of stimulus-aware video-EEG analysis, to find the correlation between the individual EEG signals and the video features (Section 2.1), while using multi-set extensions of CCA such as GCCA and CorrCA in the context of multi-subject EEG analysis, to find the correlation between EEG signals of multiple subjects watching the same video (Section 2.2). We briefly review these methods and point out the links between their seemingly-different original formulations. In Section 2.3, we explain the newly proposed video feature that we use in the video-EEG analysis of Section 2.1.

In this section, we consider the *C*-channel EEG signals **x**_*n*_(*t*)*∈* ℝ^*C*^ recorded from *N* subjects watching the same video stimulus, where *t* is the time index, and *n* is an index that refers to a specific subject (*n ∈* 1, …, *N*). The stimulus is represented by 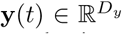, a *D*_*y*_-dimensional time-dependent feature extracted from the video, where the time index *t* is the same as the time index in the EEG signals. Without loss of generality, we assume that **x**_*n*_(*t*) and **y**(*t*) are zero-mean, i.e., 𝔼{**x**_*n*_(*t*)} = **0** and 𝔼{**y**(*t*)}= **0**. Furthermore, we assume the availability of *T* time samples, leading to the data matrices **X**_*n*_ *∈* ℝ^*T*×*C*^ and 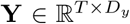, where each row is a sample of the EEG signal and the feature, respectively.

### 2.1 Stimulus-aware Video-EEG Analysis: CCA

For conciseness, we drop the subscript *n* of the EEG signals in this subsection, as the following analysis is made per subject individually. CCA was proposed in [19] as a method to find relations between two sets of variables. Given multi-dimensional EEG signal **x**(*t*) and video feature **y**(*t*), CCA computes filters **w**_*x*_ *∈ ℝ*^*C*^ (on the EEG) and 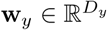 (on the stimulus) that maximize the Pearson correlation coefficient between the filtered output signals (Fig. 1a). Therefore, CCA can be formulated as the following optimization problem:

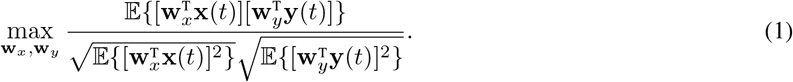

**Figure 1:**
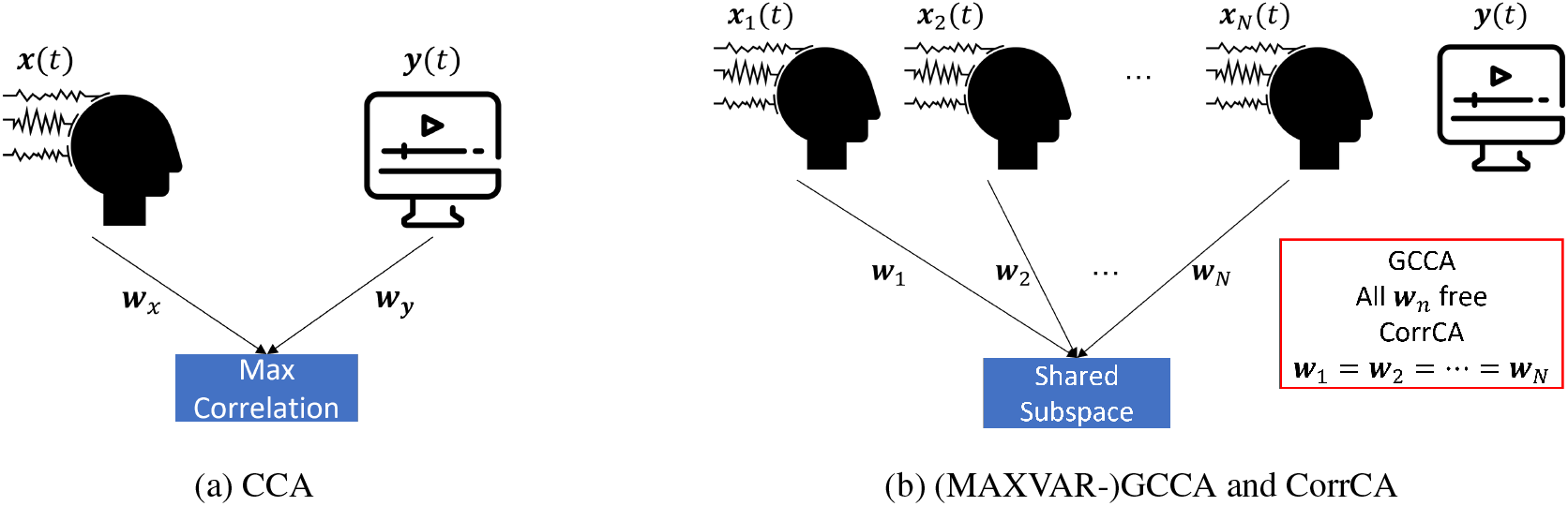
Conceptual illustrations of stimulus-aware video-EEG analysis using CCA and multi-subject EEG analysis using (MAXVAR-)GCCA and CorrCA.

The optimal **w**_*x*_ and **w**_*y*_ are called the first canonical components, and the transformed signals 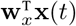 and 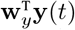 are called the first canonical directions. Here we assume the filters only act on the current timestamp, but the formulation can be easily generalized to incorporate temporal information. This can be accomplished by extending **x**(*t*) with *L*_*x*_ − 1 time-lagged copies of **x**(*t*), such that it becomes a *CL*_*x*_-dimensional vector, and extending **y**(*t*) with *L*_*y*_ − 1 time lagged copies of **y**(*t*), resulting in a *D*_*y*_*L*_*y*_-dimensional vector (similar for **w**_*x*_ and **w**_*y*_). Such extension can also compensate for the unknown time lag between the stimulus and the EEG response.

Since the scaling of **w**_*x*_ and **w**_*y*_ does not affect the objective function in (1), we can constrain the canonical directions to have unit variance to simplify the denominator. By denoting correlation matrices 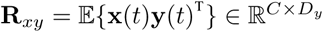, **R**_*xx*_ = 𝔼*{***x**(*t*)**x**(*t*)^T^*} ∈* ℝ^*C*×*C*^, and 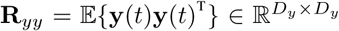 (which can be estimated as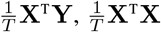 and 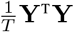, respectively), (1) can be rewritten as a constrained optimization problem:

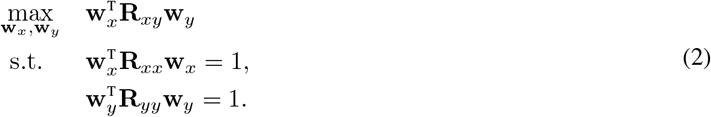

An extension of (2) in the multi-component case is:

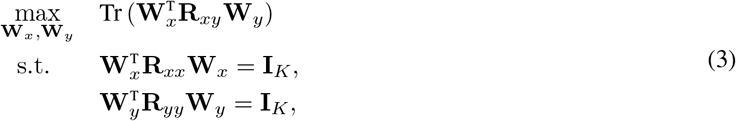

where Tr () is the trace operator, *K* is the number of components, **I**_*K*_ is the *K × K* identity matrix, and the *k*-th columns of **W**_*x*_ ∈ *ℝ* ^*C*×*K*^ and 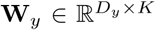 are the *k*-th canonical components. The constraints require that the canonical directions of different orders are uncorrelated to avoid trivial solutions. It can be shown that (3) can be written in a more compact form (up to a scaling factor in the solution) [20]:

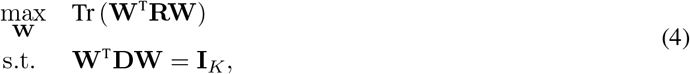

with

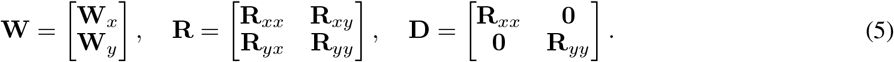

From the Karush-Kuhn-Tucker (KKT) conditions, we have the following set of equations:

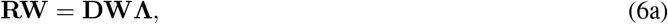

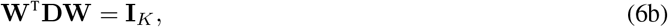

where **Λ** is a symmetric matrix containing the Lagrange multipliers. By left multiplying **W**^T^ to both sides of (6a) and making use of condition (6b), it is obvious that maximizing the objective function corresponds to maximizing Tr (**Λ**). Therefore, an optimal **W** can be obtained by solving (6a) as a generalized eigenvalue decomposition (GEVD) problem, and the columns of **W** are the generalized eigenvectors (GEVC) corresponding to the top-*K* largest generalized eigenvalues (GEVL).

### 2.2 Multi-subject EEG Analysis

#### 2.2.1 GCCA

Note that CCA only works for two views (in our case **x**(*t*) and **y**(*t*)). When analyzing the correlation between more than two views, e.g., jointly measuring the correlation between EEG signals **X**_*n*_ of multiple subjects, it becomes necessary to extend CCA to accommodate multiple matrices as inputs. The objective is now to find the per-subject filters **W**_*n*_ *∈* ℝ ^*C*×*K*^ (on the EEG) such that the outputs are on average maximally correlated. The stimulus is, in this analysis, not explicitly taken into account. However, the generalization of CCA is not unique and in this work we choose MAXVAR-GCCA [21], which aims to find linear transformations for different views such that the transformed signals, on average, closely approximate an unknown shared subspace **S** *∈* ℝ^*T*×*K*^. Mathematically, it is formulated as:

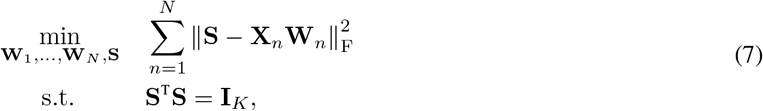

where ∥ · ∥ _F_ denotes the Frobenius norm. Using the KKT conditions, we can show that the solution to (7) is also given by a GEVD problem that has a similar form as (6a), but with a different content in the block matrices [22]:

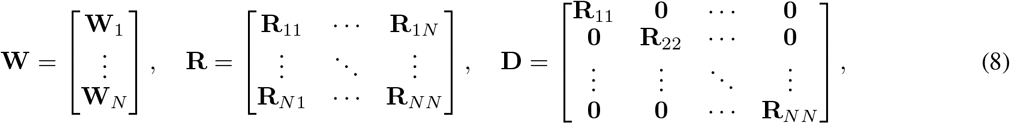

with 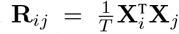, the cross-correlation matrix between the EEGs of the two subjects when *i ≠ j* and the autocorrelation matrix of subject *i* when *i* = *j*. An optimal **W** is the horizontal stack of the GEVCs corresponding to the K largest GEVLs. Therefore, up to a scaling factor in the solution, MAXVAR-GCCA is equivalent to (4) with parameters defined as in (8), which is a natural extension of CCA. The shared subspace can be obtained directly from the KKT conditions as

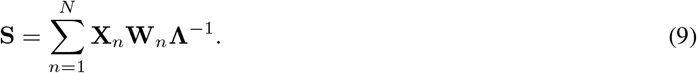

#### 2.2.2 CorrCA

CorrCA was proposed in [13], also for quantifying the correlation between the neural data of multiple subjects. A key difference between CorrCA and GCCA is that CorrCA constrains the filters of different subjects to be the same. A multi-component formulation of CorrCA is:

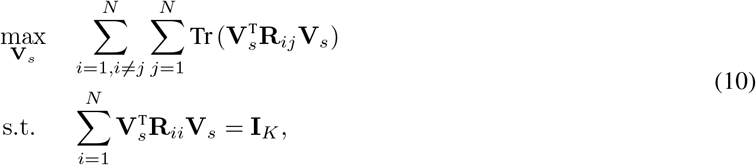

where the columns of **V**_*s*_ *∈* ℝ^*C*×*K*^ are the shared filters. From this definition, it is clear that (10) can be viewed as a straightforward multi-view extension of the (2-view) CCA in (4) with an extra constraint **W**_1_ = … = **W**_*N*_ = **V**_*s*_. One can show that CorrCA can also be viewed as a MAXVAR-GCCA with an additional constraint that all data views share the same filter (compare with (7)):

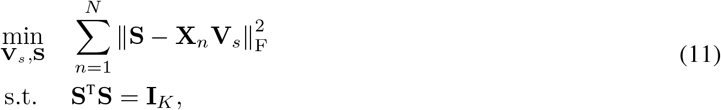

which is more insightful since it also gives a well-defined shared subspace. (10) and (11) are equivalent (up to a scaling factor), and their optimal solutions can be obtained by horizontally stacking the GEVCs corresponding to the *K* largest GEVLs of the following GEVD problem (Appendix A):

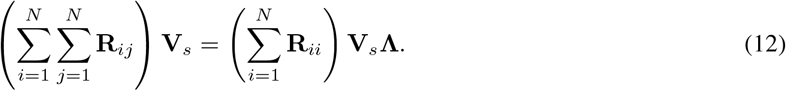

A conceptual illustration of MAXVAR-GCCA and CorrCA can be found in Fig. 1b. The constraint that the filters of different views should be the same reduces the parameters in the model and thus mitigates the overfitting problem when there is insufficient data. However, this constraint also imposes limitations on the applicability of the model, as it requires the views to have the same dimension (e.g., same number of EEG channels per subject). In comparison, GCCA allows different views to have different dimensions, so it can potentially be used to jointly analyze correlations across different data modalities or, for example, when a different number of channels or setup is used per subject. In addition, GCCA can tolerate misalignments between the data of different subjects to some extent by additionally using temporal filters. CorrCA, however, requires exact temporal alignment of the different signals for an optimal performance, even with spatial-temporal filters.

### 2.3 Video Feature Extraction

#### 2.3.1 Optical Flow and Temporal Contrast

In the limited literature on video-EEG analysis, it has been observed that optical flow and temporal contrast can be correlated with EEG signals [17]. Optical flow e stimates t he velocity vectors o f p ixels b etween consecutive frames and can thus be used to capture the motion information in videos. We applied the Gunnar-Farneback Optical Flow algorithm implemented in OpenCV to extract the flow vectors [23, 24], and computed the magnitude of each velocity vector |**v**_*m*_(**z**) |, where the subscript *m* denotes the frame index and **z** denotes the pixel coordinate. Temporal contrast is a low-level feature that is simply defined as the absolute intensity changes between consecutive frames, i.e., Δ*I*_*m*_(**z**) = |*I*_*m*_(**z**) − *I*_*m* −1_(**z**)|. In [17], both types of features are averaged across all pixels, resulting in a scalar value for each frame. We refer to the average magnitude of velocity as *AvgFlow* and the average intensity change as *AvgTempCtr*.

#### 2.3.2 Newly Proposed Object-based Features

In the previous approach of averaging over all pixels, important spatial information is lost, making it impossible to identify specific regions in a frame that elicit strong neural responses. Additionally, treating all pixels equally may not accurately reflect the attentional focus of the subject, as certain pixels, such as those corresponding to moving objects, are more likely to attract attention compared to the background, while the latter can still have a large impact on optical flow or temporal contrast. Moreover, the scaling of objects also influences the results. For example, consider an object moving at a constant speed but changing in scale. In this case, its contribution to the result varies as the number of pixels changes, which is not ideal for capturing the neural responses related to movement perception. To overcome these limitations, we propose a refined version that incorporates object segmentation. This approach involves segmenting the object(s) in each frame and calculating the mean values only for the pixels belonging to the object(s).

To perform object segmentation, we utilize Mask R-CNN [25], a deep learning model designed for object detection and segmentation in images. This model consists of two branches: one branch returns the bounding boxes of the detected objects, while the other branch provides segmentation masks. The segmentation masks are matrices where each entry indicates whether a pixel belongs to an object (1) or not (0). By feeding frames directly into the pre-trained Mask R-CNN model, we obtain the segmentation masks, which allow us to identify the pixels associated with each object. An example is shown in Fig. 2.

**Figure 2:**
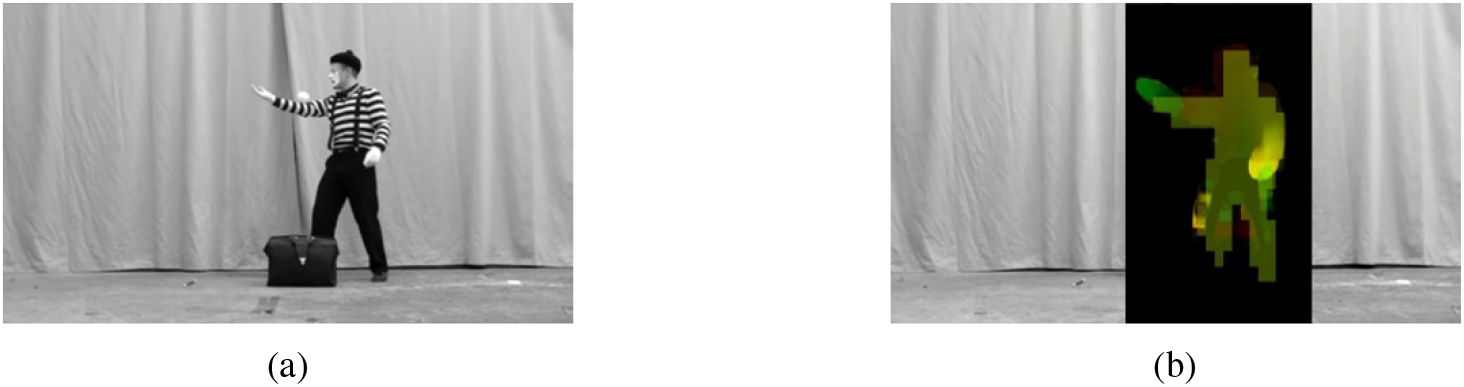
(a). A frame of a video that was used in our experiment. (b). We can obtain the bounding box and the segmentation masks using Mask R-CNN. The area within the bounding box is in black. We overlay the segmentation masks on top of it, and the color is related to the value of features.

With the obtained segmentation masks, we selectively average the flow vectors and unsigned intensity changes only over the pixels belonging to the identified object(s). In the multi-object case, one could average over the union of all pixels across all objects, or alternatively, normalize for each object separately and then sum the per-object features. A pre-selection of relevant objects could also be performed before feature fusion based on, e.g., the sizes of the bounding boxes. To circumvent feature fusion, one could treat objects separately in the analysis. Due to the many additional degrees of freedom, the multi-object case is beyond the scope of this paper and we focus on the single-object case. This object-based version of optical flow and temporal contrast is referred to as *ObjFlow* and *ObjTempCtr*, respectively. A summary of the feature definitions is given in Table 1. By incorporating object segmentation, we can retain the spatial information and stress more the regions that may receive more attention and presumably elicit higher neural responses, providing a more refined feature representation. Note that this method compensates for the variation in the number of pixels within the object since the values are averaged over pixels of the identified object.

**Table 1:**
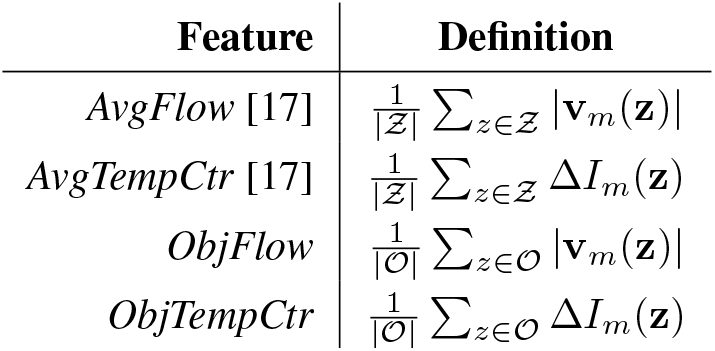
Video features and definition 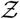 and 𝒪 denote the set of all pixels and the set of pixels belonging to the object of interest, respectively. | 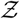 | and | *𝒪* | represent the cardinality of the sets. |**v**_*m*_(**z**)| is the magnitude of velocity.

## 3 Experiment

### 3.1 Subjects and Stimuli

20 young, healthy participants were recruited (10 females) for this study. 14 single-shot videos (duration 202 s - 463 s) showing a single moving person during various activities (dancing, mime, acrobatics, magic shows) were selected from YouTube, which were then concatenated into two longer trials (duration 36 min and 35 min) in a sequence that ensured diverse content, e.g., a dance performance followed by a mime show and a magic show, to reduce participants’ fatigue. Smooth transitions with cross-fading effects were applied between the videos. 10 of the subjects (5 females) watched an extra 24-min clip from Mr. Bean with shot cuts distributed irregularly throughout the video. For clarity, we will refer the first dataset as the Single-Shot dataset, and the second as the MrBean dataset. Unless mentioned otherwise, the same processing steps are applied to both datasets. A squared box that flashed once every 30 seconds was encoded into the videos for synchronization. It appeared outside of the original frames and was positioned in the top right corner of the screen. During passive viewing, the flashing box was covered such that it was not distractive. To avoid the confounds introduced by audio, all the videos were muted during the experiment. The study was approved by the KU Leuven Social and Societal Ethics Committee, and before participating, all participants provided their informed consent by signing a consent form.

### 3.2 Data Acquisition and Preprocessing

The EEG data of these subjects was recorded when they were watching these videos. These experiments were conducted in a quiet and dark room to minimize potential distractions. The EEG data was recorded with a BioSemi ActiveTwo system at a sample rate of 2048 Hz. The participants wore a 64-channel EEG cap, and 4 electrooculogram (EOG) sensors around the eyes were used to track eye movements. Potential bad channels were carefully logged during each session. A photo sensor for detecting the light changes of the embedded flashing box was also connected to the recorder to provide synchronization information. The collected EEG data was first segmented into different pieces corresponding to each video based on the signal of the photo sensor. Preprocessing of the EEG data was performed using functions from the MNE-Python library [26], which involved the following steps: (1) interpolating bad channels; (2) re-referencing the data to the average of all channels; (3) applying a high pass filter with a cutoff frequency of 0.5 Hz; (4) removing power line noise with a notch filter; (5) downsampling the data to 30 Hz; and (6) regressing out the EOG channels to reduce eye artifacts. For the video feature extraction, the videos were first resampled to 30 Hz and resized to 854 *×* 480 pixels. The frames were then processed as described in Section 2.3.

To avoid potential confounding effects caused by transitions between consecutive single-shot videos, the initial and final second of each video, along with the corresponding EEG data, were excluded from the analysis. Additionally, due to synchronization issues for one subject in the Single-Shot dataset, one of the single-shot videos was excluded for all subjects, resulting in a dataset of 20 subjects with 63 minutes of data each. Similarly, for the MrBean dataset, the initial and final second of the data were discarded to avoid any influence from video onset and termination. The final dataset included 10 subjects, each contributing 24 minutes of data.

### 3.3 Filter Design

In this study, we used spatial-temporal filters by default. If the data is one-dimensional, then they automatically reduce to temporal filters. For CCA, the filters applied to the video feature(s) had *L*_*y*_ = 15 lags, capturing information from the preceding 500 ms of video content. We included *L*_*x*_ = 3 lags in the filters of EEG signals, encompassing information from the previous, current, and next sample (i.e., from −33.3 ms to 33.3 ms). In GCCA or CorrCA, the filters had *L*_*x*_ = 5 lags, ranging from the past two samples to the next two samples (i.e., from −66.6 ms to 66.6 ms).

### 3.4 Evaluation Metrics

We used 10-fold cross validation to evaluate the performance of the algorithms. Each dataset was divided into 10 folds, with each fold used as the test set once while the remaining folds served as the training set. The results on the test sets were averaged across 10 folds. In the stimulus-aware video-EEG analysis, we applied the filters obtained using CCA in the training stage to the test set. For each canonical component pair *k*, we computed the Pearson correlation coefficient *ρ*_*k*_ between the transformed features and the transformed EEG signals. In addition to analyzing the results of individual canonical components, it is also valuable to evaluate the performance of multiple canonical components collectively. To achieve this, we utilized the Total Squared Correlation (TSC) metric proposed in [27]:

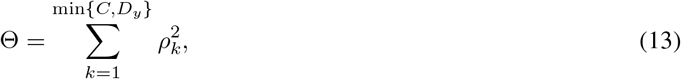

which is related to the proximity of the EEG and video feature spaces. A higher value indicates a closer correspondence. However, in this study, we were less interested in the distance between the original EEG and video feature spaces, as it may be heavily affected by noise. Therefore, we used a slightly modified version of the TSC metric to consider only the first *K* (*K* = 2 in the video-EEG analysis) canonical component pairs that were less affected by noise:

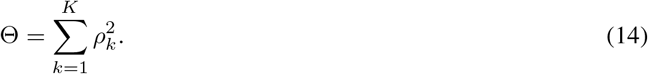

In the multi-subject EEG analysis, we applied GCCA or CorrCA filters to transform the EEG signals of each subject. We then calculated the inter-subject correlation (ISC) for each component by averaging the pairwise correlations between the transformed EEG signals of two subjects, considering all possible subject combinations [13]. Similar to the stimulus-aware video-EEG analysis, we used the TSC metric (*K* = 4) to jointly consider multiple canonical components of two subjects. By averaging the TSC values across all subject pairs, we obtained the inter-subject TSC (ISTSC).

To assess the significance of the results, we employed a permutation test. The null hypothesis assumed that the transformed data views were uncorrelated. To simulate this scenario, we first obtained the transformed data with the trained filters, and then created the permuted copies by randomly shuffling the transformed EEG samples (or/and the transformed features) 1000 times. The p-value was determined as the proportion of absolute correlation coefficients (or ISC values) calculated from the permuted data that exceeded the correlation between the original transformed data. If the p-value was below the *α*-level (set to 0.05), we rejected the null hypothesis and concluded that the correlation was statistically significant.

For comparing the performance of different features, we employed the paired Wilcoxon signed-rank test, a non-parametric statistical hypothesis test for comparing paired observations. In our study, pairs could be, e.g., a pair of TSCs obtained with two different features for a subject. The null hypothesis assumed that the median difference between the pairs of observations was zero. In two-sided tests, the alternative hypothesis was that the median difference was not equal to zero. We primarily used one-sided tests in this study, where the alternative hypothesis was that the median difference was greater (lower) than zero. We applied a Bonferroni correction when multiple comparisons were performed.

The quality of the features can also be evaluated based on their performance in a specific task. A common practice involves using these features in a match-mismatch (MM) task [9, 28]. An illustration of the MM task is shown in Fig. 3. In this study, we trained the filters of the EEG signals and video features using CCA, and divided the test set into *N*_*s*_ 1-min EEG and video segments. For each EEG segment in the test set, there were (*N*_*s*_ − 1) test pairs, where each pair consisted of the matching video segment combined with a non-matching segment. We applied filters obtained in the training stage and computed the correlations between the transformed EEG segment and the two transformed video segments, respectively. The video segment with a higher correlation was selected as the match. The total number of tests conducted was *N*_*s*_(*N*_*s*_ − 1), and the error rate was calculated as the proportion of incorrect decisions. We also analyzed the dual problem, i.e., for each video segment, we created EEG segment pairs and performed the MM task. The error rate was averaged across 10 folds.

**Figure 3:**
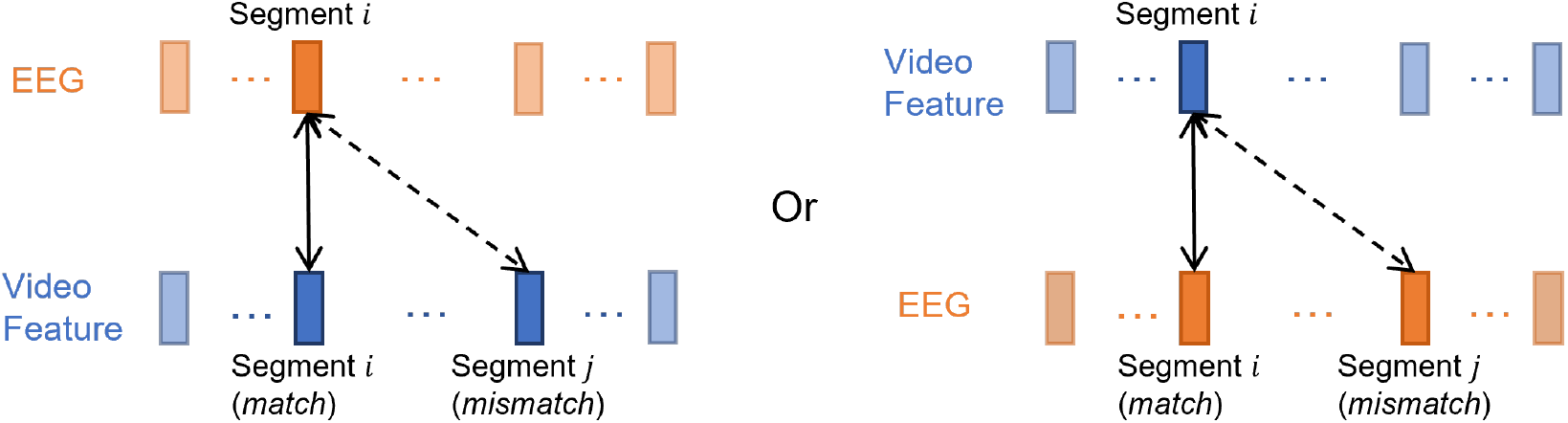
An illustration of the match-mismatch task. Given an EEG (video) segment, create segment pairs containing the matching and a non-matching video (EEG) segment. For each segment pair, decide which video (EEG) segment matches the EEG (video) segment.

### 3.5 Interpretation of Weights

Correlation coefficients and TSCs provide valuable insights into the degree of correlation between data views or the proximity of data spaces. However, to gain further understanding, we can also look into the weights of the filters. For temporal filters, a conventional practice to interpret the weights is to plot the frequency response, which shows the frequency components that are amplified or attenuated by the filter. Regarding spatial filters, an intuitive way is to identify the channels with the highest contribution by looking at the absolute values of the weights and then locate the regions of interest. However, as argued in [29], this approach can be misleading since channels that are not highly relevant to the extracted components may also receive large weights due to, e.g., noise cancelling. A better approach is to compute the forward models, which involves reconstructing the original EEG signals from the extracted components, and then plot the topographic maps of the weights of the forward models (for both the video-EEG and multi-subject EEG analysis). The extracted components are better reflected in channels with larger (unsigned) weights. As the unrelated EEG background noise can, in theory, not be predicted from these components, these forward models purely reflect the stimulus-related contributions and not the noise.

For CCA, following the notations in Section 2.1 and adding subscripts to matrices to specify the subject, we define the transformed individual EEG signal as **S**_*n*_ = **X**_*n*_**W**_*n*_ *∈* ℝ^*T*×*K*^ and the individual forward model as **F**_*n*_ *∈* ℝ^*C*×*K*^ for subject *n*. The forward model can be computed by solving the following LS problem:

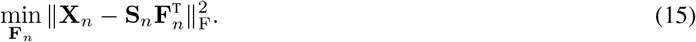

The solution is

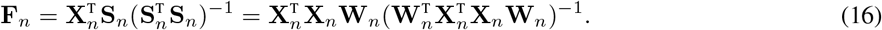

For GCCA (and CorrCA), we reconstruct signals from the shared subspace **S** and define an averaged version of the forward model **F** as

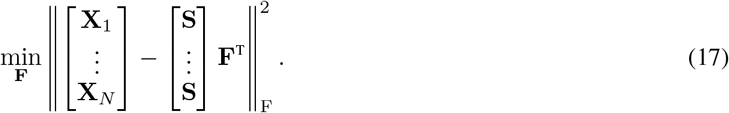

Making use of the constraint that **S**^T^**S** = **I**_*K*_ and a KKT condition 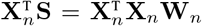 of (7), the solution can be obtained as:

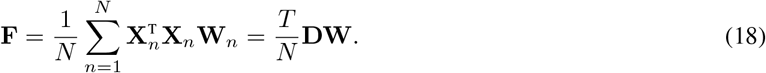

Note that when using spatial-temporal filters, the form of the forward models will be slightly different from (16) and (18) since **S** (and **S**_*n*_) will be computed from a lagged version of **X**_*n*_. For each component, we can generate a topographic plot illustrating the weights of the forward model. To ensure clarity in visualization, we always plot the absolute values of the weights.

## 4 Results

### 4.1 Mind shot cuts

Using the MrBean dataset, we first show that the shot cuts in the videos lead to prominent peaks in *AvgFlow* and *AvgTempCtr*, which actually dominate the correlations found by CCA.

An underlying assumption in Gunnar-Farneback Optical Flow (Section 2.3.1) is that the objects in a video sequence tend to move coherently and smoothly, which is clearly violated when there is a shot cut. Most of the pixels in the previous frame will have large displacement or even have disappeared in the next frame, resulting in a spurious peak in *AvgFlow* (Fig. 4a). Similarly, shot cuts usually lead to large intensity changes, resulting in similar peaks in *AvgTempCtr* (Fig. 4c). To automatically identify these peaks, we used a peak detection algorithm (find_peaks [30]) on the *AvgFlow* (or *AvgTempCtr*) feature, yielding 124 (or 125) prominent peaks. To verify that these peaks correspond to shot cuts, we applied a shot change detection method (AdaptiveDetector() from the PySceneDetect library [31]) to the video, which returned 127 time points of the shot cuts. Out of the 124 peaks in *AvgFlow* (and 125 peaks in *AvgTempCtr*), 122 (and 124, respectively) were matched with the shot cuts.

**Figure 4:**
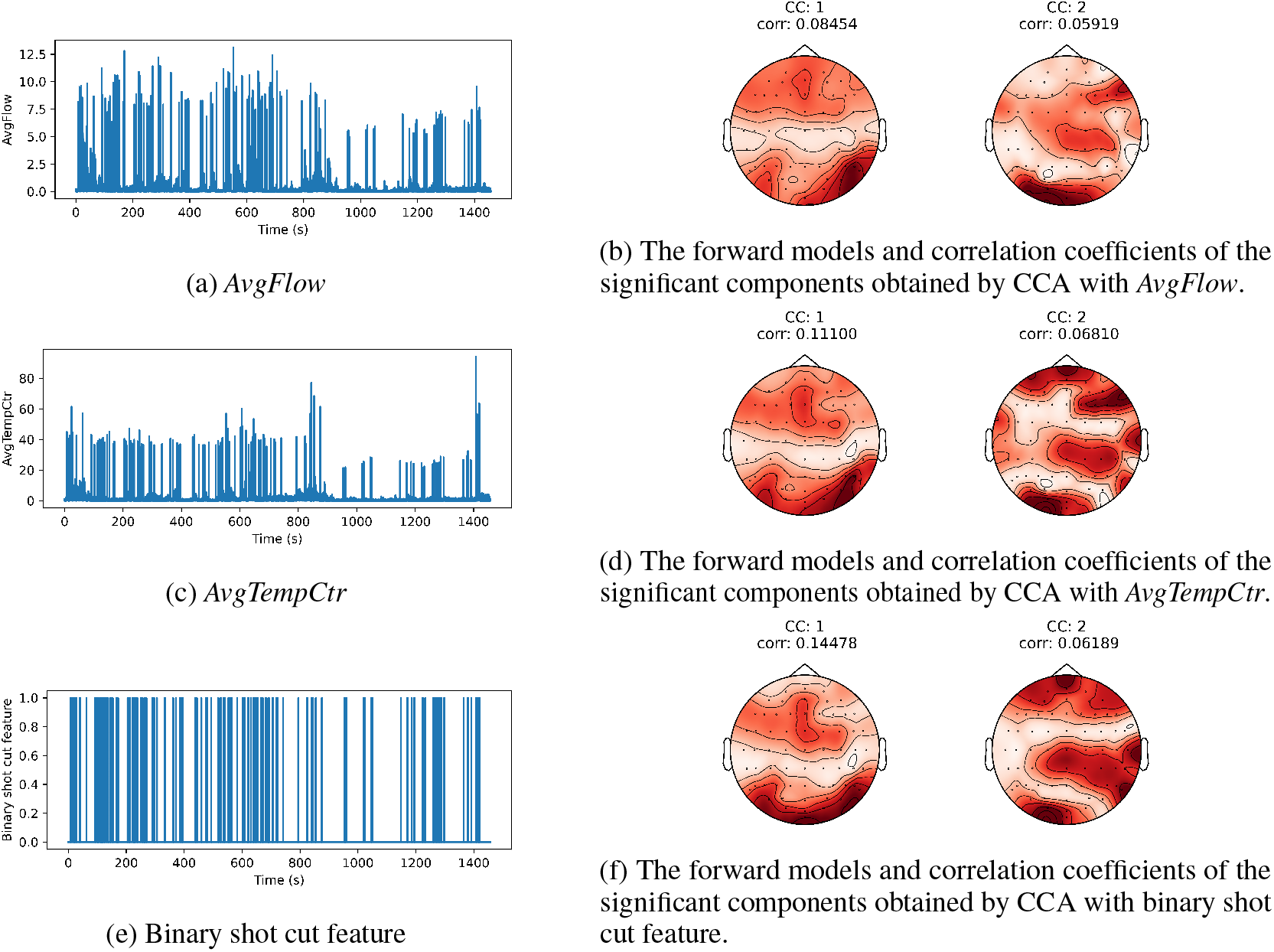
Comparison between performing CCA between the EEG and video stimulus using *AvgFlow, AvgTempCtr*, and the binary shot cut feature for a representative subject. The plotted canonical components are all significant. With the binary shot cut feature, the correlation coefficients are higher or comparable to those obtained by CCA with *AvgFlow* and *AvgTempCtr*. The forward models also share similar patterns when using the three different features.

Since the amplitude of the peaks caused by shot cuts no longer indicates the magnitude of velocity or the temporal intensity change, these peaks should be treated as artifacts. Therefore, apart from applying CCA to the original features, we explored a second setting where we first identified shot cuts based on the results of the shot change detection function. Subsequently, we removed one second of data before and after each shot cut, and performed CCA again. This procedure resulted in a loss of approximately 4 minutes of data, which accounted for less than 20% of the entire dataset, and allows to probe the correlations with *AvgFlow* and *AvgTempCtr* when no shot cuts are present. Thirdly, instead of removing shot cuts, it is also interesting to see what would happen if we exclusively retain shot cut information. For this purpose, we designed a binary shot cut feature that assigned a value of 1 if a shot cut was present in the corresponding frame, and 0 otherwise (Fig. 4e). In this third setting, we used CCA to correlate the binary shot cut feature with EEG signals. The results of the first two canonical components under all three settings are shown in Table 2.

**Table 2:**
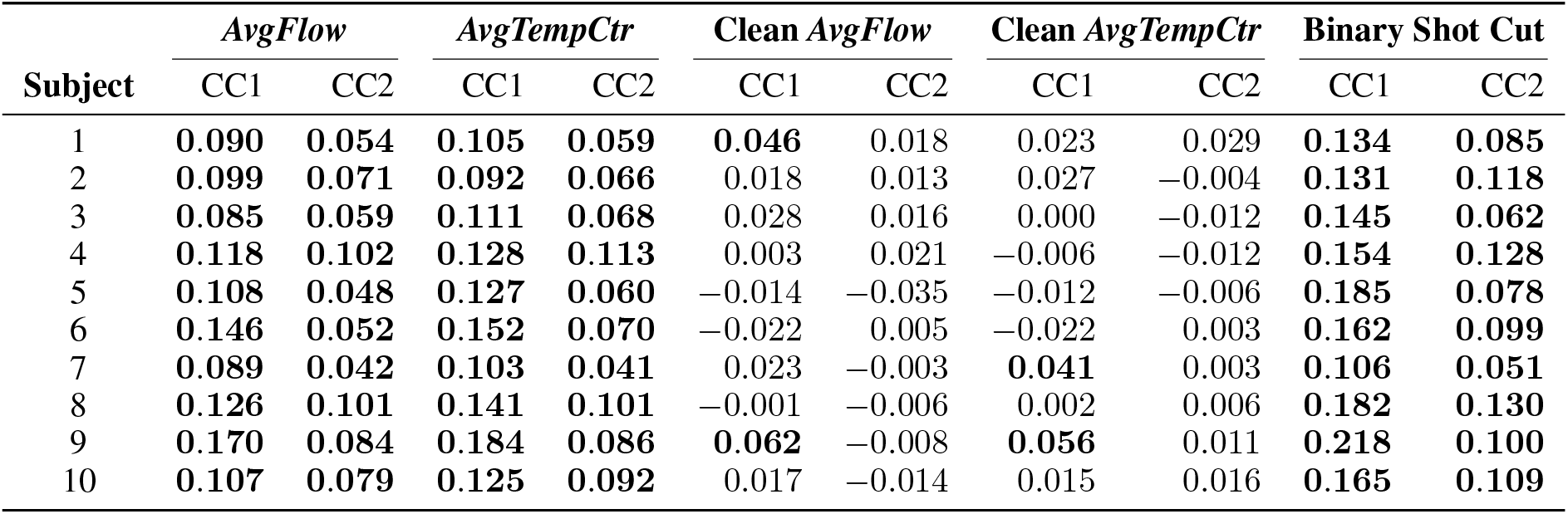
Pearson correlation coefficients of the first two canonical components (CC) obtained on the MrBean dataset with CCA under three settings: 1) *AvgFlow* (*AvgTempCtr*): correlating original *AvgFlow* (*AvgTempCtr*) features with EEG signals; 2) Clean *AvgFlow* (Clean *AvgTempCtr*): correlating *AvgFlow* (*AvgTempCtr*) features with EEG signals when shot cuts are removed (both from EEG and the features); 3) Binary Shot Cut: correlating binary shot cut features with the EEG signals. The significant correlations are in bold.

By comparing the results of the first two settings, we can see that the correlations drop drastically and even become non-significant after removing the shot cuts, both for *AvgFlow* and *AvgTempCtr*. In contrast, the correlations obtained with the binary shot cut feature are comparable to or even higher than the ones obtained with the original *AvgFlow* or *AvgTempCtr* feature. We can also observe in Fig. 4b, 4d, and 4f (topographic plots for a representative subject) that the forward models of the canonical components obtained with *AvgFlow, AvgTempCtr*, and the binary shot cut feature are similar to each other. Based on these observations, we argue that the correlations obtained with *AvgFlow* and *AvgTempCtr* using CCA are primarily driven by shot cuts. When shot cuts are removed from the natural videos, these features are inadequate in consistently finding significant correlations, suggesting their limited utility in capturing the temporal coupling between EEG signals and natural single-shot videos.

Since the binary shot cut feature exhibits a strong correlation with EEG signals, we infer that shot cuts elicit robust (probably ERP-like) neural responses in the brain. Given that the data of all subjects are synchronized, it is expected that the EEG components associated with shot cuts also show coherence across subjects and can thus be captured by GCCA/CorrCA. After removing the shot cuts, those EEG components may again disappear or change. To test this intuition, we applied CorrCA to the EEG data, both with and without shot cuts removed. CorrCA was chosen as it adds additional regularization, which is required given the relatively small scale (24 min *×* 10 subjects) of the dataset. Fig. 5 illustrates the correlations and forward models of the top-10 canonical components before and after the removal of shot cuts. The number of significant components decreased from 10 to 6. The vanished components may be associated with the neural responses elicited by shot cuts, which also implies that one factor can affect multiple EEG components. Among the remaining significant components, the correlations decrease, and the reduction may also be related to shot cuts.

**Figure 5:**
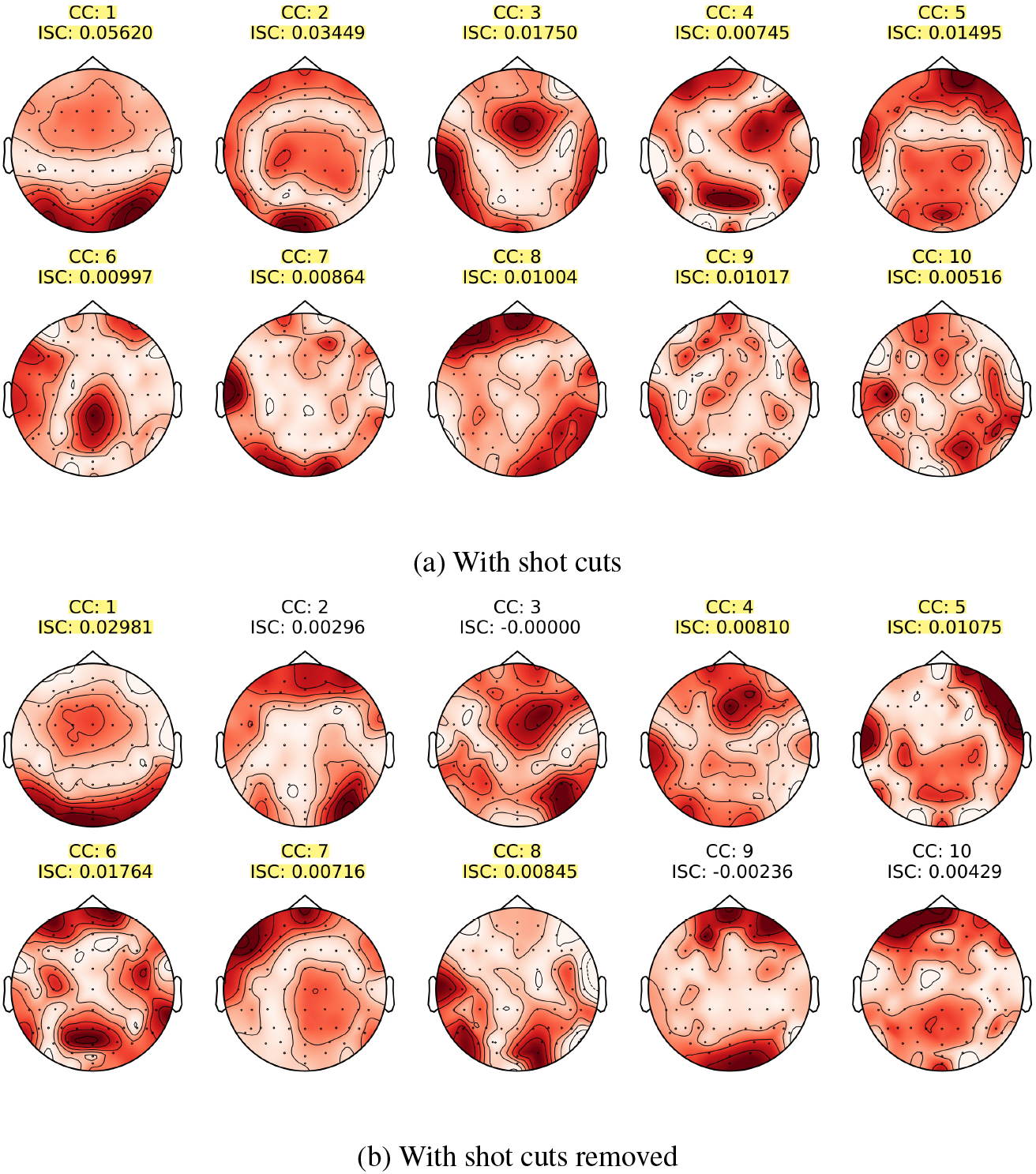
Forward models and ISCs obtained by CorrCA with the EEG data of 10 subjects watching Mr. Bean with and without shot cuts. The significant components are highlighted.

These results demonstrate that shot cuts have a significant impact on both video-EEG analysis with the traditional *AvgFlow* and *AvgTempCtr* features and multi-subject EEG analysis. These shot cuts lead to significant correlations, which are useful as they could indicate whether a subject is watching a video or not. However, in many application scenarios, visual stimuli typically do not contain shot cuts. For instance, when determining a driver’s focus on road conditions using EEG signals and video streaming from a dashcam, consecutive frames will have smooth transitions. Moreover, Table 2 clearly shows that *AvgFlow* and *AvgTempCtr* are unable to extract significant components in-between shot cuts, such that more temporally fine-grained estimations of levels of attention or engagement are impossible. To confirm these intial findings, in the next section, we will investigate the performance of *AvgFlow* and *AvgTempCtr* in single-shot videos, without any shot cuts, and propose our novel object-based features from Section 2.3.2 as alternatives.

### 4.2 Object-based Features Lead to Significant Correlation in Single-shot Videos

Starting from this section, we analyze the Single-Shot dataset, with video clips without shot cuts. In Table 3, we compare the performance of CCA using the original features *AvgFlow* and *AvgTempCtr* with their newly proposed object-based counterparts, *ObjFlow* and *ObjTempCtr*. It is evident that the use of object-based features leads to substantial improvements in robustness, i.e., the feature is able to consistently find significant correlations in all subjects. In terms of *AvgFlow*, only 1 subject exhibits two significant components, while only 25% of the correlations across both components and all subjects is significant. In comparison, by employing the proposed *ObjFlow*, we are able to identify at least one significant component for each subject, with 85% of the correlations significant across both components and all subjects. While *AvgTempCtr* demonstrates slightly better robustness compared to *AvgFlow*, incorporating *ObjTempCtr* again clearly results in a drastic improvement.

**Table 3:**
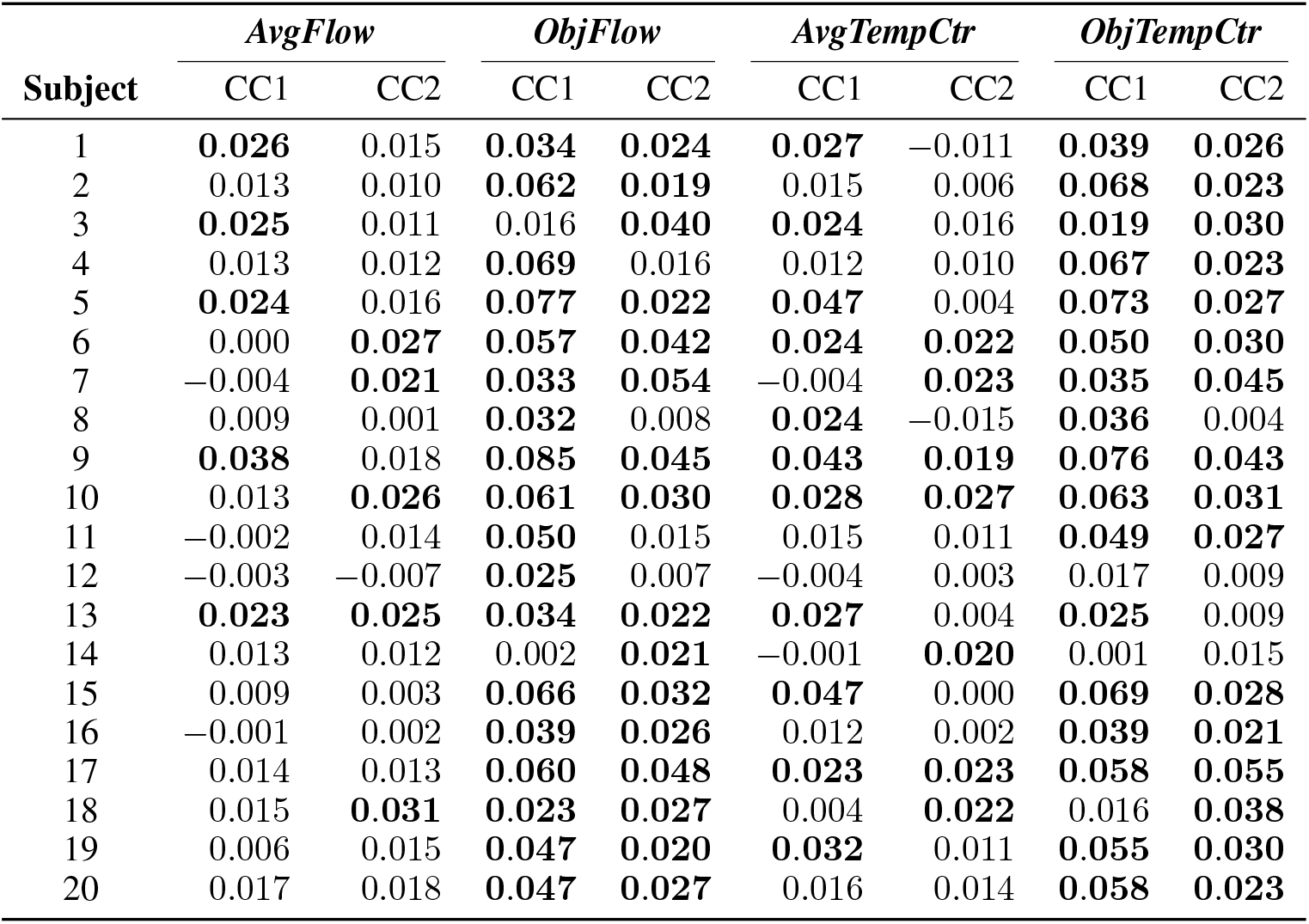
Pearson correlation coefficients of the first two canonical components (CC) obtained from CCA using *AvgFlow, AvgTempCtr*, and their object-based versions *ObjFlow* and *ObjTempCtr*. The significant correlations are in bold.

| Subject | *AvgFlow* |  | *ObjFlow* |  | *AvgTempCtr* |  | *ObjTempCtr* |  |
| --- | --- | --- | --- | --- | --- | --- | --- | --- |
|  | CC1 | CC2 | CC1 | CC2 | CC1 | CC2 | CC1 | CC2 |
| 1 | **0.026** | 0.015 | **0.034** | **0.024** | **0.027** | −0.011 | **0.039** | **0.026** |
| 2 | 0.013 | 0.010 | **0.062** | **0.019** | 0.015 | 0.006 | **0.068** | **0.023** |
| 3 | **0.025** | 0.011 | 0.016 | **0.040** | **0.024** | 0.016 | **0.019** | **0.030** |
| 4 | 0.013 | 0.012 | **0.069** | 0.016 | 0.012 | 0.010 | **0.067** | **0.023** |
| 5 | **0.024** | 0.016 | **0.077** | **0.022** | **0.047** | 0.004 | **0.073** | **0.027** |
| 6 | 0.000 | **0.027** | **0.057** | **0.042** | **0.024** | **0.022** | **0.050** | **0.030** |
| 7 | −0.004 | **0.021** | **0.033** | **0.054** | −0.004 | **0.023** | **0.035** | **0.045** |
| 8 | 0.009 | 0.001 | **0.032** | 0.008 | **0.024** | −0.015 | **0.036** | 0.004 |
| 9 | **0.038** | 0.018 | **0.085** | **0.045** | **0.043** | **0.019** | **0.076** | **0.043** |
| 10 | 0.013 | **0.026** | **0.061** | **0.030** | **0.028** | **0.027** | **0.063** | **0.031** |
| 11 | −0.002 | 0.014 | **0.050** | 0.015 | 0.015 | 0.011 | **0.049** | **0.027** |
| 12 | −0.003 | −0.007 | **0.025** | 0.007 | −0.004 | 0.003 | 0.017 | 0.009 |
| 13 | **0.023** | **0.025** | **0.034** | **0.022** | **0.027** | 0.004 | **0.025** | 0.009 |
| 14 | 0.013 | 0.012 | 0.002 | **0.021** | −0.001 | **0.020** | 0.001 | 0.015 |
| 15 | 0.009 | 0.003 | **0.066** | **0.032** | **0.047** | 0.000 | **0.069** | **0.028** |
| 16 | −0.001 | 0.002 | **0.039** | **0.026** | 0.012 | 0.002 | **0.039** | **0.021** |
| 17 | 0.014 | 0.013 | **0.060** | **0.048** | **0.023** | **0.023** | **0.058** | **0.055** |
| 18 | 0.015 | **0.031** | **0.023** | **0.027** | 0.004 | **0.022** | 0.016 | **0.038** |
| 19 | 0.006 | 0.015 | **0.047** | **0.020** | **0.032** | 0.011 | **0.055** | **0.030** |
| 20 | 0.017 | 0.018 | **0.047** | **0.027** | 0.016 | 0.014 | **0.058** | **0.023** |

Apart from comparing the robustness across subjects, we can also directly compare the strength of correlations. From Table 3, we observe that the first canonical components do not always capture the most correlated information in practice, although they should theoretically. Indeed, sometimes the second component exhibits a higher correlation, which may be attributed to overfitting on the training set or small differences between the training and test set. Due to these inconsistencies in the ordering of the components (according to correlation values) between the training and test set, it makes more sense to consider the canonical components jointly using TSC. The results are plotted in Fig. 6. To compare the performance of different features, we conducted one-sided Wilcoxon signed-rank tests. The results indicate that *ObjFlow* and *ObjTempCtr* exhibit significant superiority over *AvgFlow* and *AvgTempCtr*, with p-values *<* 0.001 in both cases.

**Figure 6:**
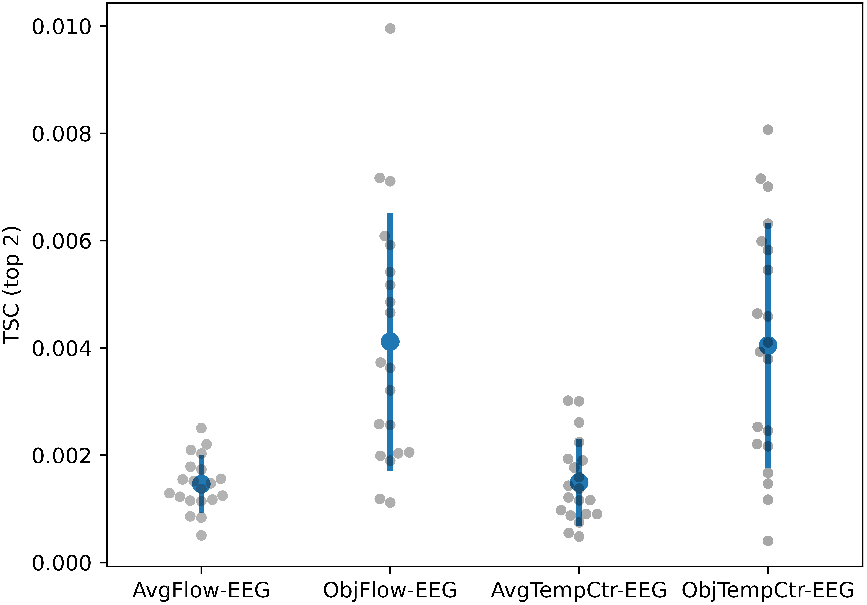
The per-subject individual TSCs (denoted by grey dots) between the EEG signal and *AvgFlow, ObjFlow, AvgTempCtr*, and *ObjTempCtr*. The bars indicate the mean and standard deviation of the TSCs across subjects.

These results confirm the results from Section 4.1, i.e., that the traditional *AvgFlow* and *AvgTempCtr* are inadequate to capture the temporal correlations between the EEG and an attended video when there are no shotcuts. Moreover, the results clearly show that *ObjFlow* and *ObjTempCtr* are better features that consistently yield significant correlations. This agrees with our intuition that the feature is more specific and emphasizes the regions where viewers are likely to focus. However, it is worth noting that *ObjFlow* may be correlated with eye movements if participants tracked the object during passive viewing. Consequently, the residual eye motion artefacts in the EEG signals could potentially drive significant correlations (despite regressing out the EOG signals from the EEG data), posing challenges in determining whether higher-level cognitive processes, such as movement perception, contribute to these correlations. A similar issue may arise with *ObjTempCtr*, as it implicitly encodes object motion information. Therefore, we will further investigate whether the eye movements drive these correlations in the following section.

### 4.3 Are Correlations Driven by Eye Movements?

To check whether the correlations are driven by eye movements, we correlated the EOG signals with the proposed object-based features and compared the results with those obtained from EEG signals in the Single-Shot data set. We observed a significant decrease in TSCs of *ObjFlow*-EOG and *ObjTempCtr*-EOG (Fig. 7), with p-values *<* 0.001. This suggests that the leakage of EOG signals into the EEG cannot fully explain the correlations between the EEG signals and our features; instead, neural activities captured by EEG dominate the correlations. We conclude that eye movements do not dominantly drive the correlations.

**Figure 7:**
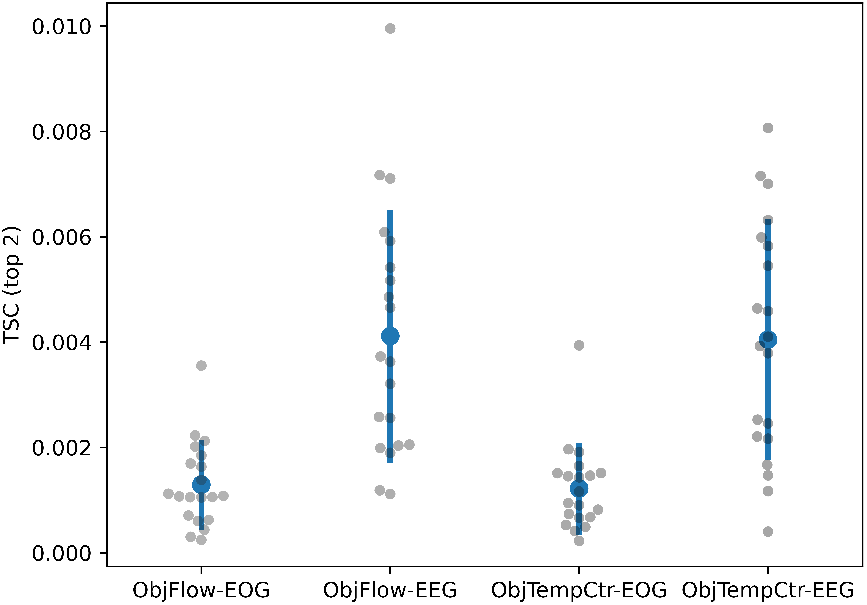
The per-subject individual TSCs (denoted by grey dots) between the EOG (EEG) signal and proposed object-based features. The bars indicate the mean and standard deviation of the TSCs across subjects.

### 4.4 Object-based Features Perform Better in MM Task

In Section 4.2, we have demonstrated that object-based features exhibit higher correlations with EEG signals compared to traditional features. To further demonstrate the superiority of these object-based features in a more quantitatively verifiable setting, we conducted the MM task on the Single-Shot dataset using *AvgFlow, ObjFlow, AvgTempCtr*, and *ObjTempCtr*, respectively, as described in Section 3.4. We retained the top 5 canonical components obtained with CCA in the training set. In the decision-making phase, we selected the one with highest correlation for each EEG-video pair, and from each pair we selected the segment with the highest correlation as the “matched” segment. The error rates are presented in Fig. 8. In both scenarios, i.e., matching video segments given EEG segments and matching EEG segments given video segments, object-based features yielded lower error rates compared to their traditional counterparts, with p-values *<* 0.001. This further supports the argument that the proposed object-based features are more effective in identifying meaningful temporal correlations between EEG and video signals.

**Figure 8:**
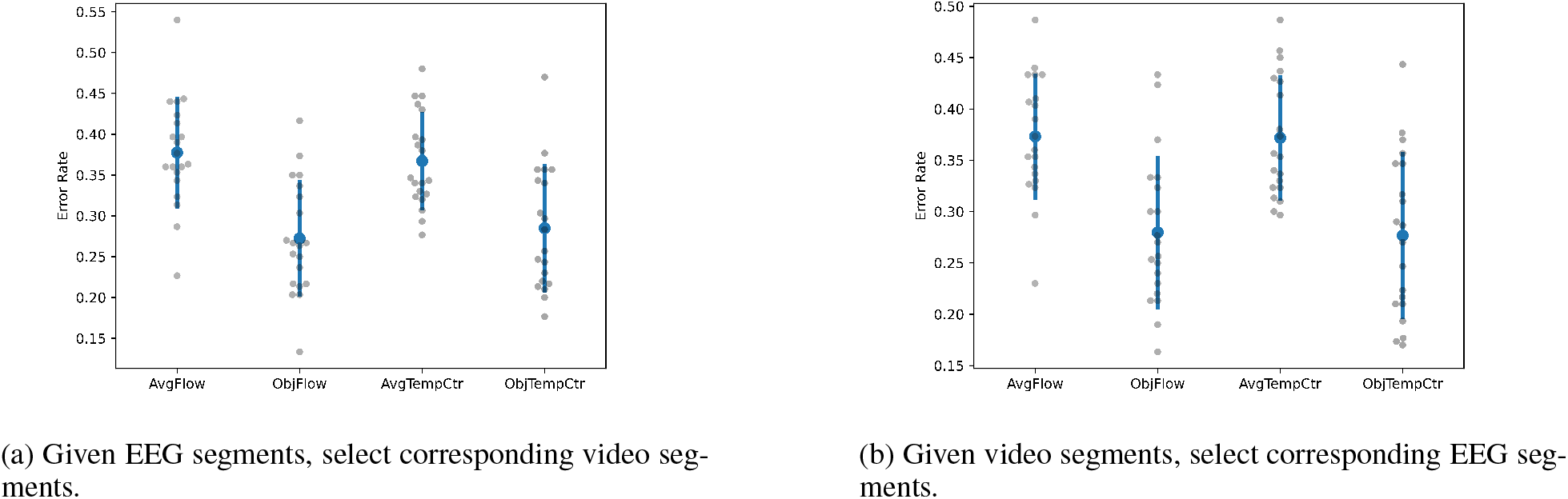
The per-subject individual error rates of the MM task (denoted by grey dots) using *AvgFlow, ObjFlow, AvgTempCtr*, and *ObjTempCtr*. The bars indicate the mean and standard deviation of the error rates across subjects.

### 4.5 Multi-subject EEG Analysis on Single-shot Videos

One limitation of doing stimuli-response analysis using CCA is that it can only be performed individually and thus cannot leverage information across subjects. With multi-subject EEG analysis, we can extract the shared subspace of EEG signals of all the subjects, which is spanned by the coherent EEG components that are time-locked to the video stimuli. These coherent EEG components have higher SNR since the asynchronous noise and background EEG activities are suppressed. Therefore, it is informative to show their forward models, which can reveal the regions where these coherent EEG components are more reflected. Given that the Single-Shot dataset is a larger dataset (63 min *×* 20 subjects) than the MrBean dataset (24 min *×* 10 subjects) used in Section 4.1, we chose GCCA instead of CorrCA for the multi-subject analysis. The ISCs and the forward models (defined in (17)) of the first 10 canonical components are shown in Fig. 9, 8 of which are significant. The ISTSC of the top 4 canonical components is 0.0066.

**Figure 9:**
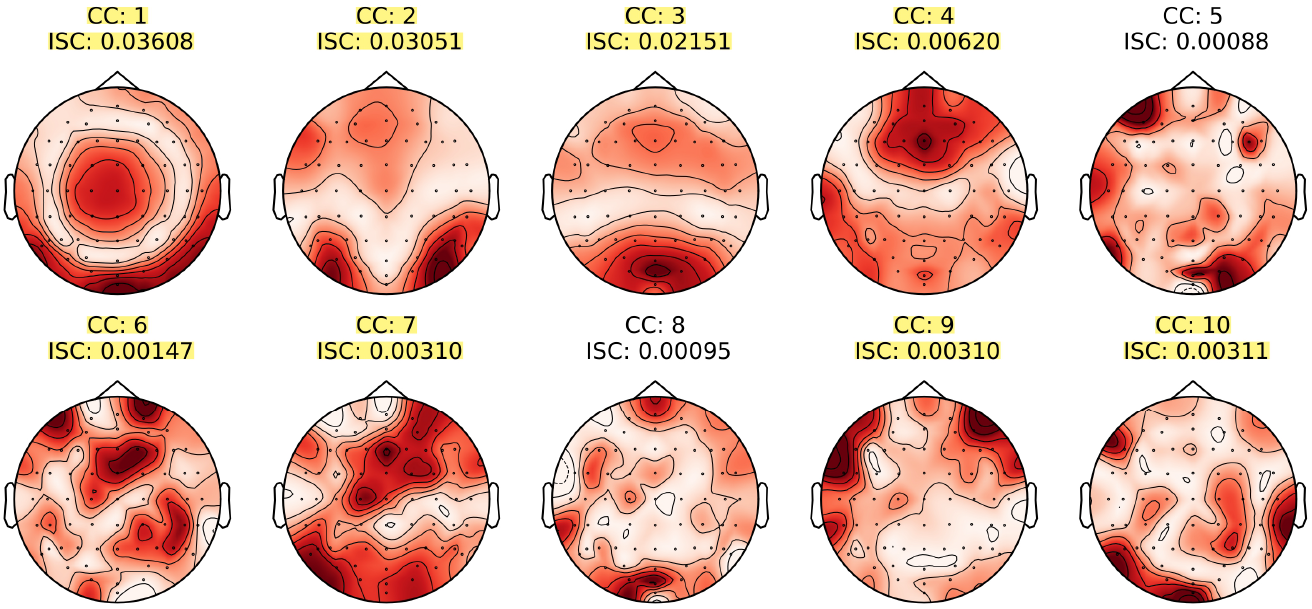
ISCs and forward models of top 10 EEG components obtained by GCCA. The significant components are highlighted.

### 4.6 Proportion of Variance Explained

A follow-up question arising from the multi-subject EEG analysis is whether we can isolate a subset of the obtained coherent EEG components that are dominantly driven by our features. This can be achieved qualitatively by first regressing out the features from the EEG signals (for the entire dataset), then reapplying the GCCA algorithm and identifying the components that either disappear or exhibit substantial changes. Results of GCCA with *ObjFlow* regressed out from EEG signals can be found in Fig. 10. Notably, these results closely resemble those in Fig. 9, particularly for the first 4 components with higher ISCs, suggesting that *ObjFlow* may not be the dominant feature for any of them. Since *ObjTempCtr* is highly correlated with *ObjFlow* in our dataset (indicated by a correlation coefficient of 0.857), the forward models when *ObjTempCtr* is regressed out are similar and we omit them for brevity.

**Figure 10:**
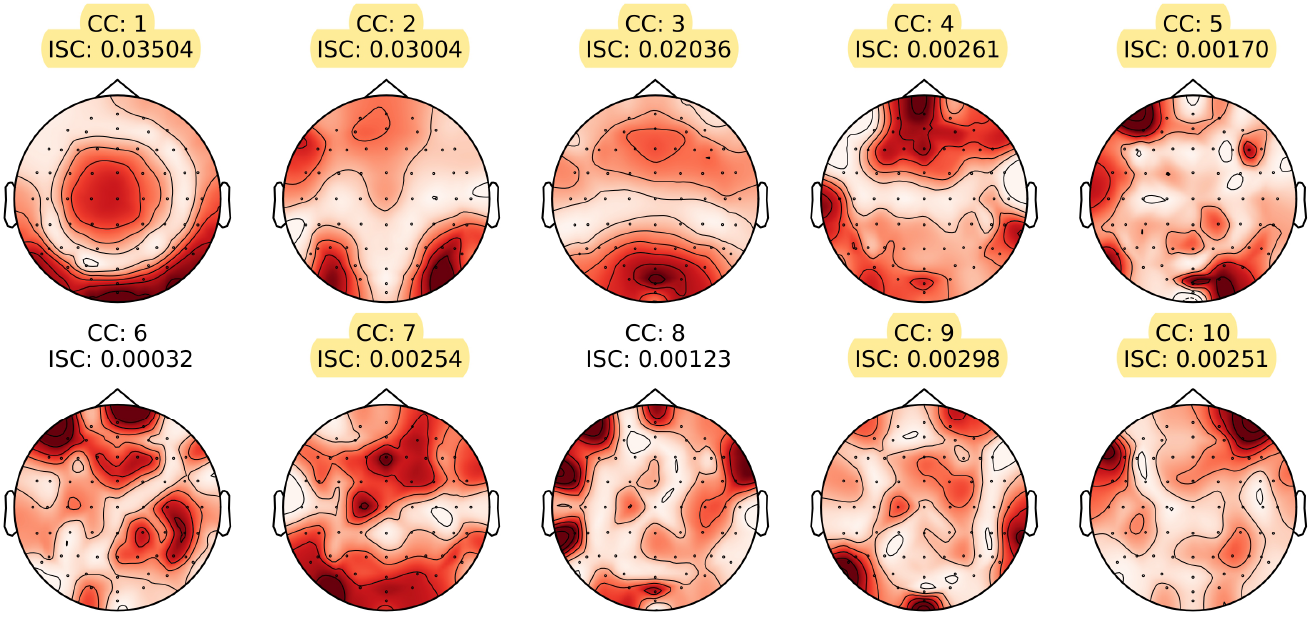
ISCs and forward models of top 10 EEG components obtained by GCCA with *ObjFlow* regressed out from the EEG signals. The significant components are highlighted.

To quantitatively assess the extent to which coherent EEG components obtained by GCCA can be attributed to our features, we can calculate the proportion of variance in the coherent stimulus responses explained by *ObjFlow* (or *ObjTempCtr*). Specifically, the variance of the stimulus responses in the *k*-th coherent EEG component can be estimated by the averaged pairwise covariance of the transformed EEG signals, i.e., 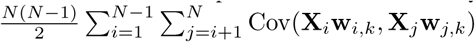 where **w**_*i,k*_ denotes the *k*-th column of **W**_*i*_. We refer to this quantity as inter-subject covariance (ISCOV). Since the (incoherent) background EEG components are orthogonal across subjects, they will not influence this ISCOV, except for a residue due to finite sample sizes. In order to estimate the error on the ISCOV due to this residue, we performed the following bootstrap procedure: randomly shifting the data in the test set across subjects with at least 10 s, using the pre-trained GCCA filters to transform the shifted data, and then computing the ISCOV. This procedure was repeated 100 times, with the outcomes aggregated across all components. Since the ISCOV is an averaged value across different subject pairs, the obtained ISCOVs can be modeled as a Gaussian distribution according to the central limit theorem. The 95% confidence interval of the ISCOV, obtained using the shifted EEG data and serving as the error estimate, is (−4.4 *×* 10^−9^, 4.3 *×* 10^−9^).

Table 4 presents the ISCOVs when using the original EEG data and using the EEG data with *ObjFlow* or *ObjTempCtr* regressed out. From the numbers, it appears that component 4 is most related to our features given that the ISCOV decreases by 58.7% with *ObjFlow* regressed out and by 63.5% with *ObjTempCtr* regressed out. It is also notable that the ISCOVs no longer exceed the upper limit of the 95% confidence interval for the estimation error after regression. However, despite these reductions, the forward model of component 4 does not exhibit significant changes, which might imply that our features could be coincidentally highly correlated with the actual features driving the responses in component 4.

**Table 4:** ISCOVs (1 *×* 10^−8^) of the first 10 canonical components (CC) obtained by GCCA under three conditions: using (1) the original EEG data; (2) EEG data with *ObjFlow* regressed out; (3) EEG data with *ObjTempCtr* regressed out. The 95% confidence interval for the estimation error is (−4.4 *×* 10^−9^, 4.3 *×* 10^−9^).

|  | CC1 | CC2 | CC3 | CC4 | CC5 | CC6 | CC7 | CC8 | CC9 | CC10 |
| --- | --- | --- | --- | --- | --- | --- | --- | --- | --- | --- |
| Original | 4.32 | 3.64 | 2.33 | 0.63 | 0.09 | 0.13 | 0.31 | 0.10 | 0.28 | 0.33 |
| *ObjFlow* regressed out | 4.16 | 3.57 | 2.18 | 0.26 | 0.17 | 0.02 | 0.25 | 0.10 | 0.28 | 0.24 |
| *ObjTempCtr* regressed out | 4.11 | 3.56 | 2.21 | 0.23 | 0.16 | 0.03 | 0.22 | 0.11 | 0.28 | 0.24 |

To aggregate the effects across different components, we define the total variance as the sum of ISCOVs over the selected components. The proportion of variance explained by *ObjFlow* (or *ObjTempCtr*) is then computed as the complement of the ratio between the total variance obtained using the EEG data with *ObjFlow* (or *ObjTempCtr*) regressed out and that using the original EEG data, i.e.,

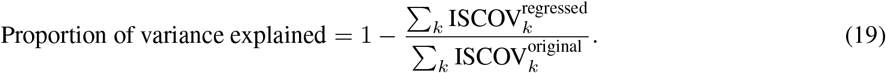

For the top 4 most prominent coherent components whose ISCOVs exceed the upper limit of the 95% confidence interval for the estimation error before regression, the proportion of variance explained is 6.9% by *ObjFlow* and 7.4% by *ObjTempCtr*. In other words, around 93% of the variance in the coherent stimulus responses remains unexplained, which indicates the presence of potentially more dominant yet undiscovered features.

## 5 Discussions

### 5.1 Shot Cuts Highly Influence Temporal Correlations

In [32], the authors observed that the ISC based on fMRI recordings is different during the viewing of unedited and edited videos of dance performance. The unedited version represented a continuous view captured from a single camera, while the edited version consisted of concatenated shots from different cameras, therefore included shot cuts. The authors calculated ISC maps for each video and found that the two individual maps (edited vs unedited) exhibited broad overlap, but the edited version showed more significant voxels. These findings align with our own results in Section 4.1, where we discovered that in the multi-subject analysis, the correlations were stronger when shot cuts were present and decreased significantly when shot cuts were removed. Additionally, we also observed substantial overlap in the forward models of the first components when comparing the results with or without shotcuts (Fig. 5).

In a recent study by Nentwich et al. [33], participants were presented with natural videos while neural responses were recorded using intacranical EEG (also known as electrocorticography) with 6328 electrodes implanted throughout the entire brain. The analysis was performed channel-wise on an average brain: they extracted the broad-band high-frequency amplitude (BHA) ranging from 70 to 150 Hz and modeled the BHA of each channel as a convolution of the visual stimulus and an unknown TRF, which can be estimated using LS. While the primary focus of their research was on investigating the effects of semantic changes, there was a finding relevant to our study: they observed that a greater number of channels responded to film cuts compared to visual motion calculated using optical flow. This finding supports our conclusion that shot cuts dominate correlations in video-EEG analysis especially when using the traditional, non-object based versions of the optical flow and temporal contrast.

In earlier EEG studies with natural video footage as stimuli such as [13, 16, 17], the shot cuts were not removed and their effect was not the focus. Therefore, it is likely that the correlations found in these studies were almost exclusively driven by shot cuts. In [16], it was noted that the highest and the most sustained ISC coincided with the video segment having the most scene changes in the example video clip, which aligns with our argument. While in certain cases, e.g., using the correlation as a marker of engagement [13], the origins of the correlations are less important, it is still advisable to be aware of the impact of shot cuts since they elicit strong neural responses that could potentially overshadow more intricate responses related to higher-level cognitive processes.

### 5.2 Interpretation and Relations of Optical Flow and Temporal Contrast

In our experiments, both *ObjFlow* and *ObjTempCtr* lead to higher correlations with the EEG signals and lower error rates in the MM task. However, there is no significant difference in the performance of these two features. The p-value of the two-sided paired Wilcoxon signed-rank test on the TSCs obtained using *ObjFlow* and *ObjTempCtr* yields 0.90. The error rates when matching video segments given EEG segments and matching EEG segments given video segments are also not significantly different, with p-values of 0.14 and 0.59, respectively. Notably, the forward models exhibit remarkable similarity for these two features, as shown in Fig. 11 for a representative subject. Furthermore, stacking the two features together as a two-dimensional time series and inputting it into CCA does not show any significant difference either. These outcomes are probably attributed to the high correlation between *ObjFlow* and *ObjTempCtr* within our video dataset, as indicated by a correlation coefficient of 0.857. The high correlation comes as a surprise since *ObjFlow* and *ObjTempCtr* seem to be unrelated, representing motion and intensity changes, respectively. However, one can show that the two features are implicitly coupled, posing challenges in distinguishing their individual effects on the EEG signals (Appendix B).

**Figure 11:**
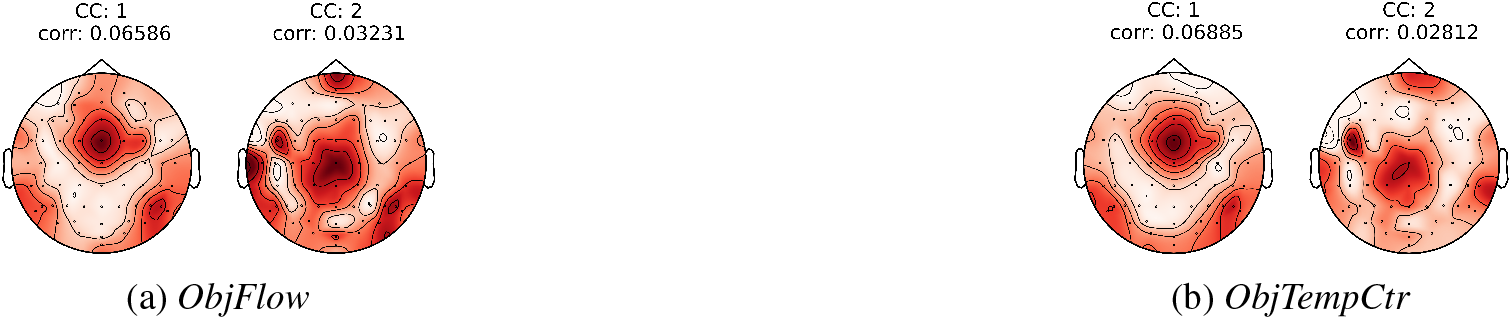
Comparison of forward models obtained using CCA with *ObjFlow* and *ObjTempCtr* for a representative subject. For these two significant components, the forward models are highly similar.

Therefore, based solely on the results obtained from the current dataset, we cannot conclusively determine the driving factor behind the observed correlations. While this issue is not the primary focus of our paper, we highlight it to emphasize the non-trivial nature of identifying the underlying causes of correlations when using natural videos. One feature could encode information of features with very different physical meanings, and it is difficult to disentangle their effects using uncontrolled visual stimuli. Additionally, if one feature works well in certain videos but not in others, it suggests that the feature may coincidentally correlate with the true feature that elicits the neural responses. Therefore, the diversity of videos in the training set is important for the correct interpretation of features.

### 5.3 Interpretation of Neural Patterns

Research on human movement perception also provides valuable insights into the interpretation of our results, particularly regarding the activation patterns observed in certain brain areas. For example, Grosbras et al. conducted a meta-analysis combining fMRI results from multiple studies focused on three categories of motion: face, hands, and whole body movements [34]. They applied the activation likelihood estimation method with random effect analysis to generate a probability map reflecting the likelihood that a particular voxel was activated. Their findings revealed convergence of brain activation in the occipito-temporal and fronto-parietal regions across all categories, although with different peak locations and extents. In [35], dancers were asked to perform specific movements while detailed movement features were gathered using accelerometers. Videos of these dancers were shown to fMRI participants, and the collected data were analyzed. The researchers discovered that low-level features such as acceleration corresponded to brain regions associated with early visual and motion-sensitive areas. On the other hand, mid-level features such as dynamic symmetry mapped to the occipito-temporal cortex, posterior superior temporal sulcus, and superior parietal lobe. These findings could provide an explanation on why, in our forward models obtained with ObjFlow using CCA (e.g., Fig. 11a), the occipito-temporal and fronto-parietal regions exhibited higher levels of activation compared to other areas. However, as discussed in Section 5.2, the activation of certain regions may also be related to the perception of brightness. It was found in [36] that the neural responses in the striate cortex explicitly encode brightness changes, which could be an additional explanation for the activation of the occipital area.

### 5.4 Potential Usage of Object-based Features

Object-based features provide more refined representations, leading to higher and more reliable correlations with the EEG signals. These correlations could be employed as a metric for overall attention levels or engagement [18]. Additionally, object-based features may open the door for attention decoding, identifying which object the participants are focusing on. In [37], the results indicated that participants’ selective attention mechanisms operated efficiently, being able to isolate and focus on the object of their choice despite the presence of multiple objects embedded within complex backgrounds in each scene. Therefore, by correlating features extracted from different objects with the EEG signals using the video-EEG analysis with the newly proposed features, it may be possible to determine the object of interest and detect shifts in visual attention. An expected constraint is that the signatures of different objects should be sufficiently distinct, otherwise the results for different objects may be too similar. Apart from attention decoding, it is also desirable to measure the overall level of attention, potentially also in a multi-subject setting. However, fusing the features extracted from different objects is a challenging problem that requires further investigation.

### 5.5 Quest for Novel Video Features

Observing Fig. 5, we noted a decrease in ISCs and the number of significant components when shot cuts were removed from the MrBean dataset. Nevertheless, it is encouraging that there are still significant correlations remaining, indicating that the neural responses related to shot cuts are not the sole factors coherent across subjects. The same holds true for the Single-Shot dataset, where multiple significant components were found despite the absence of shot cuts in the videos (Fig. 9). This motivates the quest for novel video features that are not solely driven by shot cuts and can capture these components. The object-based features we have proposed are limited by their constraint on the number of objects in each frame. Furthermore, these features explain only approximately 7% of the variance in the coherent stimulus responses across subjects (Section 4.6), suggesting that there are potentially more dominant features that are yet to be discovered. This is not particularly surprising, as both optical flow and temporal contrast are still relatively low-level features. It could be beneficial to leverage knowledge from computer vision and representation learning to investigate higher-level and more abstract features. Such features could potentially prove useful across a wide range of videos.

## 6 Conclusion

This study focused on identifying temporal correlations between natural video footage and EEG signals, for which two new datasets were collected. The MrBean dataset used a film clip that contains many shot cuts as the stimulus, while in the Single-Shot dataset the videos were carefully selected to be shot cut-free and to contain only a single moving object. We revealed that the correlations between video features such as optical flow and temporal contrast and the EEG signals, which were reported in previous studies, were heavily influenced by shot cuts present in the videos, leading to over-optimistic correlations between both modalities. We showed that removal of such shot cuts result in non-significant correlations in the majority of the subjects. We proposed the use of object-based features as a more robust alternative, resulting in significant correlations with the EEG signals across all subjects, even in the absence of shot cuts. Importantly, we showed that the observed correlations were not predominantly driven by eye movements, which are usually considered as confounds. Furthermore, we demonstrated that the proposed object-based features were more effective in the MM task, yielding lower error rates compared to traditional features. Finally, we illustrated that the proposed features did not dominantly drive the coherent stimulus responses, and more influential features are yet to be discovered.

For future research, there are several promising directions worth exploring. Firstly, we can shift from linear models to non-linear models to capture more complex relationships between video features and EEG signals. Secondly, exploring higher-level video features, whether with or without semantic meanings, may provide insights on how the brain processes more abstract information. Lastly, applying the proposed object-based features in a multi-object setting would reveal their effectiveness in decoding attention towards specific objects within complex visual scenes.

## Acknowledgment

This research is funded by the Research Foundation - Flanders (FWO) project No G081722N, junior postdoctoral fellowship fundamental research of the FWO (for S. Geirnaert, No. 1242524N), the European Research Council (ERC) under the European Union’s Horizon 2020 research and innovation program (grant agreement No 802895), the Flemish Government (AI Research Program), and the PDM mandate from KU Leuven (for S. Geirnaert, No PDMT1/22/009). All authors are also affiliated with Leuven.AI - KU Leuven institute for AI, Belgium.

The authors also thank the participants for their time and effort in the experiments. The code and the datasets will be made publicly available upon publication.

## Appendix A Equivalence of (10) and (11)

In this section, we show that the solutions of (10) and (11) are the same up to a scaling factor. We start with the solution of (10). The Lagrange function of (10) is:

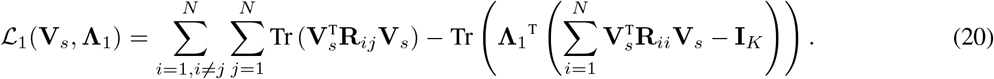

The KKT conditions for **V**_*s*_ to be optimal are then given by:

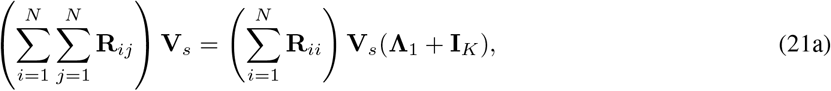

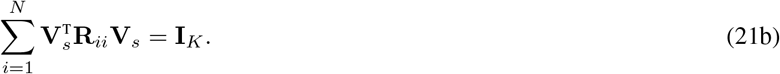

By left multiplying (21a) by **V**_*s*_^T^ and using (21b), the objective function of (10) can be simplified as Tr (Λ_1_). Therefore, the optimal **V**_*s*_ is the horizontal concatenation of the GEVCs corresponding to the *K* largest GEVLs of the GEVD problem (21a) (up to orthogonal transformations). The correct scaling of the GEVCs is determined by (21b). Similarly, for (11), we write down the Lagrangian:

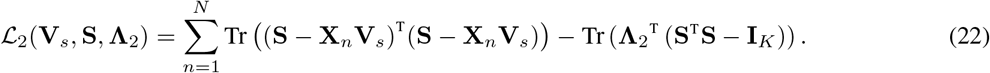

The KKT conditions for optimal **V**_*s*_ and **S** are:

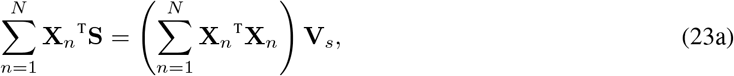

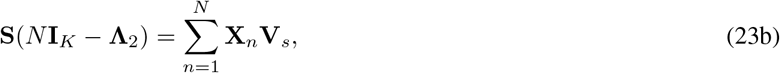

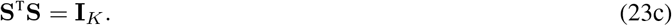

Plugging (23b) into (23a) yields:

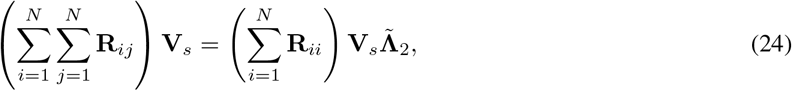

with 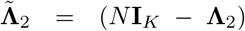. Using (23a) and (23c), the objective function of (11) can be written as 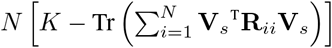. From (23b) and (23c), we have 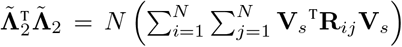. Combining it with (24) yields 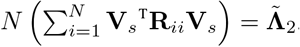. Therefore, minimizing the objective function is equivalent to maximizing 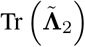. Then, again, the optimal **V**_*s*_ is the horizontal stack of the GEVCs corresponding to the *K* largest GEVLs of the GEVD problem (24). The scaling factor is determined by (23c). As (21a) and (24) represent the same GEVD problems, the solutions of (10) and (11) are identical up to a scaling factor.

## Appendix B The Coupling Between Optical Flow and Temporal Contrast

To illustrate the implicit coupling between *ObjFlow* and *ObjTempCtr*, we start with the calculation of velocity vectors in optical flow, which usually involves making assumptions on the pixel intensity. Take the Gunnar-Farneback Optical Flow [23] as an example. The algorithm is based on polynomial expansions, approximating the intensity *I*_*m*_(**z**) of some neighborhood of each pixel in the *m*-th frame with, e.g., a local quadratic polynomial *I*_*m*_(**z**) = **z**^T^**A**_*m*_(**z**)**z** + **b**_*m*_(**z**)^T^**z** + *c*_*m*_(**z**), where **z** denotes the two-dimensional pixel coordinate. The local parameters{**A**_*m*_(**z**), **b**_*m*_(**z**), *c*_*m*_(**z**)} of this model for frame *m* can be estimated using weighted LS. The algorithm then tries to find the displacement vector of each pixel **d**_*m*_(**z**) under the assumption that *I*_*m*_(**z**) = *I*_*m* − 1_(**z** − **d**_*m*_(**z**)). By matching the coefficients of the two polynomials, we can derive **d**_*m*_(**z**) as

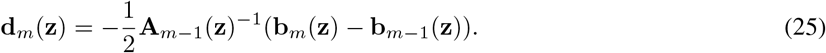

The velocity vector **v**_*m*_(**z**) can then be obtained by multiplying **d**_*m*_(**z**) with the sampling frequency of the video *f*_*s*_. Additionally, the coefficient matching yields the following two equations: **A**_*m*_(**z**) = **A**_*m* −1_(**z**) and *c*_*m*_(**z**) = *c*_*m* −1_(**z**) + **d**_*m*_(**z**)^T^**A**_*m*−1_(**z**)**d**_*m*_(**z**) **b**_*m*−1_(**z**)^T^**d**_*m*_(**z**). Since in practice **A**_*m*_(**z**) = **A**_*m*−1_(**z**) generally does not hold, the approximation [**A**_*m*_(**z**) + **A**_*m*−1_(**z**)]*/*2 is usually used for both **A**_*m*_(**z**) and **A**_*m*−1_(**z**). If we follow the assumptions of Gunnar-Farneback Optical Flow and utilize the derived equations, the temporal contrast can be expressed in terms of the displacement vector and the coefficients of the quadratic polynomial:

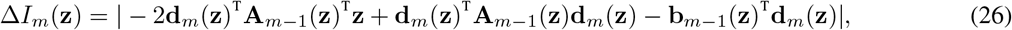

which clearly shows the interdependence between optical flow and temporal contrast.

## Notes

### Competing Interest Statement

The authors have declared no competing interest.

